# RB dependent transcriptional regulation at mitotic centromeres preserves genome stability

**DOI:** 10.1101/2025.06.04.657825

**Authors:** Elizabeth A Crowley, Amity L Manning

## Abstract

Transcripts derived from centromere repeats play a critical role in the localization and activity of kinetochore components during mitosis such that disruption of RNA polymerase II-dependent transcription compromises the fidelity of chromosome segregation. Here, we show that the retinoblastoma tumor suppressor protein (RB), a critical regulator of the G1/S cell cycle transition, additionally plays an important role in the regulation of centromere transcription during mitosis. We find that cells lacking RB experience increased RNA Polymerase II activity at mitotic centromeres and a corresponding increase in nascent RNA transcripts derived from centromere sequences. Together with high levels of centromere transcription and corresponding R-loop formation, RB-deficient cells exhibit centromere DNA breaks and local activation of ATR that correspond with increased centromere localization of Aurora B, destabilization of kinetochore-microtubule attachments, and an increase in anaphase defects. Importantly, reduction of DNA damage, ATR activity, and mitotic defects following inhibition of RNA Pol II, or targeted repression of centromere transcription through centromere tethering of Suv420H2, support that mitotic defects in RB-deficient cells are linked to centromere transcription.

## Introduction

During cell division newly replicated chromosomes are segregated equally into two daughter cells, ensuring the faithful inheritance of genetic information. This process is dependent on a specialized region of the chromosome known as the centromere (Brinkley & Stubblefield, 1966; Musacchio & Salmon, 2007) which is epigenetically defined by the presence of nucleosomes containing the histone variant Centromere Protein A (CENP-A) (Earnshaw *et al*, 1986; Fachinetti *et al*, 2013; Palmer *et al*, 1991). Proteinaceous structures known as kinetochores are then assembled at each centromere where they govern the interaction between chromosomes and microtubules of the mitotic spindle.

Long thought to be transcriptionally inert, we now appreciate that both the core centromere region and the pericentric flanking regions are actively transcribed by RNA polymerase II (RNAPII) (Chan *et al*, 2012; Ideue *et al*, 2014; Rošić & Erhardt, 2016; Wong *et al*, 2007). Transcriptional activity at centromeres is a conserved property among species (Melters *et al*, 2013; Talbert *et al*, 2018) and has an important role in regulating centromere and kinetochore composition (reviewed in (Perea-Resa & Blower, 2018)). A key function of centromere transcription is to promote chromatin-remodeling that is permissive for loading of CENP-A-containing nucleosomes during the G_1_ phase of the cell cycle (Bobkov *et al*, 2018; Dunleavy *et al*, 2011; Jansen *et al*, 2007).

However, the transcription of centromeres is not limited to interphase cells and RNAPII is active at mitotic centromeres (Chan *et al*., 2012). During cell division, centromere transcripts act as molecular tethers that can recruit and activate key centromere and kinetochore proteins, including CENP-C (Bury *et al*, 2020; McNulty *et al*, 2017; Politi *et al*, 2002) which functions to stabilize CENP-A retention (Falk *et al*, 2015; Guo *et al*, 2017; McNulty *et al*., 2017; Watanabe *et al*, 2019), AurB kinase, a member of the chromosomal passenger complex (CPC) that modulates kinetochore-microtubule attachments (Blower, 2016; Ferri *et al*, 2009; Jambhekar *et al*, 2014; Quénet & Dalal, 2014), and Shugoshin 1, which regulates centromere cohesion (Chen *et al*, 2021; Liu *et al*, 2015). The formation of transcription-dependent R loops, where ssDNA is displaced as the transcription machinery moves along the template strand, also recruits the ataxia-telangiectasia and Rad3-related protein kinase (ATR) (Matos *et al*, 2020). During mitosis, ATR functions, in part, to active Aurora B kinase (AurB) locally at the centromere (Kabeche *et al*, 2018). Together, ATR and AurB dynamically regulate kinetochore microtubules to promote accurate chromosome segregation during cell division.

The variety of ways in which transcriptional activity at the centromere contributes to mitotic fidelity indicates that genetic or epigenetic changes that alter the accessibility of the centromere to transcription factors or otherwise perturb the level of centromere transcription during mitosis, have the potential to impact both kinetochore composition and the fidelity of chromosome segregation. Consistent with this model, two functionally relevant marks of transcriptionally silent heterochromatin, trimethylation of lysine 9 on Histone H3 (H3K9me3) and trimethylation of lysine 20 on Histone H4 (H4K20me3), are highly enriched at pericentromeric regions and yet are restricted from centromeres (Agredo & Kasinski, 2023; Schotta *et al*, 2004). Experimental manipulations that permit the spread of heterochromatin into the centromere corrupt deposition of the centromere specific CENPA- containing nucleosomes and lead to chromosome segregation errors (Martins *et al*, 2020; Sidhwani & Straight, 2023). Interestingly, the absence or loss of heterochromatin from the pericentromere can also compromise the accuracy of chromosome segregation (Herlihy *et al*, 2021; Martins *et al*, 2016; Martins *et al*., 2020).

The retinoblastoma tumor suppressor protein, RB, physically interacts with the enzymes that place the heterochromatic marks H3K9me3 (Suv39) and H4K20me3 (Suv420h2) (Sanidas *et al*, 2019). RB-dependent recruitment of Suv420h2 is relevant for establishment of pericentric H4K20me3 (Gonzalo & Blasco, 2005; Gonzalo *et al*, 2005) and loss of RB results in decreased H4K20me3 and cohesin complex (a reader of H4K20me3) at pericentromeres (Gonzalo *et al*, 2005(Manning *et al*, 2010; Manning *et al*, 2014). Here we demonstrate that high levels of centromere transcription and corresponding mis-regulation of mitotic kinases underlie centromere damage and mitotic errors that result from loss of RB. Furthermore, we show that suppression of mitotic transcription, centromere-targeted restoration of epigenetic silencing, or titration of kinase activity are sufficient to restore mitotic fidelity in cells lacking the RB tumor suppressor. Together these findings indicate that epigenetic regulation of centromeres is a dynamic and targetable process by which to modulate the accuracy of chromosome segregation.

## Results

### RB loss promotes RNA Polymerase II-dependent centromere transcription

To determine whether loss of RB impacts transcription during mitosis, we first examined cells with and without RB depletion for evidence of RNA polymerase II (RNAPII) activity. Using human telomerase reverse transcriptase (hTERT)- immortalized retinal pigment epithelial cells (RPE-1) engineered to carry an inducible RB-targeting shRNA construct (hTERT-RPE-1 shRB), we treated cells with 2 µg/mL of doxycycline for 48 hours to induce RB depletion (Figure 1A). Nocodazole arrested mitotic cells were collected and chromosome spreads were prepared for immunofluorescence analysis of total RNAPII and active RNAPII (phosphorylated at Serine 2, which is indicative of elongating polymerase) (Figure 1B). RNAPII activity has previously been described at mitotic centromeres (Chan et al, 2012). Consistent with these reports, hTERT-RPE-1 cells were positive for both total (RNAPII) and active (RNAPII pS2) RNA polymerase II. Using quantitative measures of RNAPII and RNAPII pS2 signal intensity across anti-centromere-antigen (ACA)-labelled kinetochores, we observe that while depletion of RB does not affect the total pool of RNAPII at mitotic centromeres, it does lead to a greater than 4-fold increase in RNAPII pS2, indicating that cells lacking RB experience an increase in transcriptional activity at mitotic centromeres (Figure 1C).

**Figure 1:**
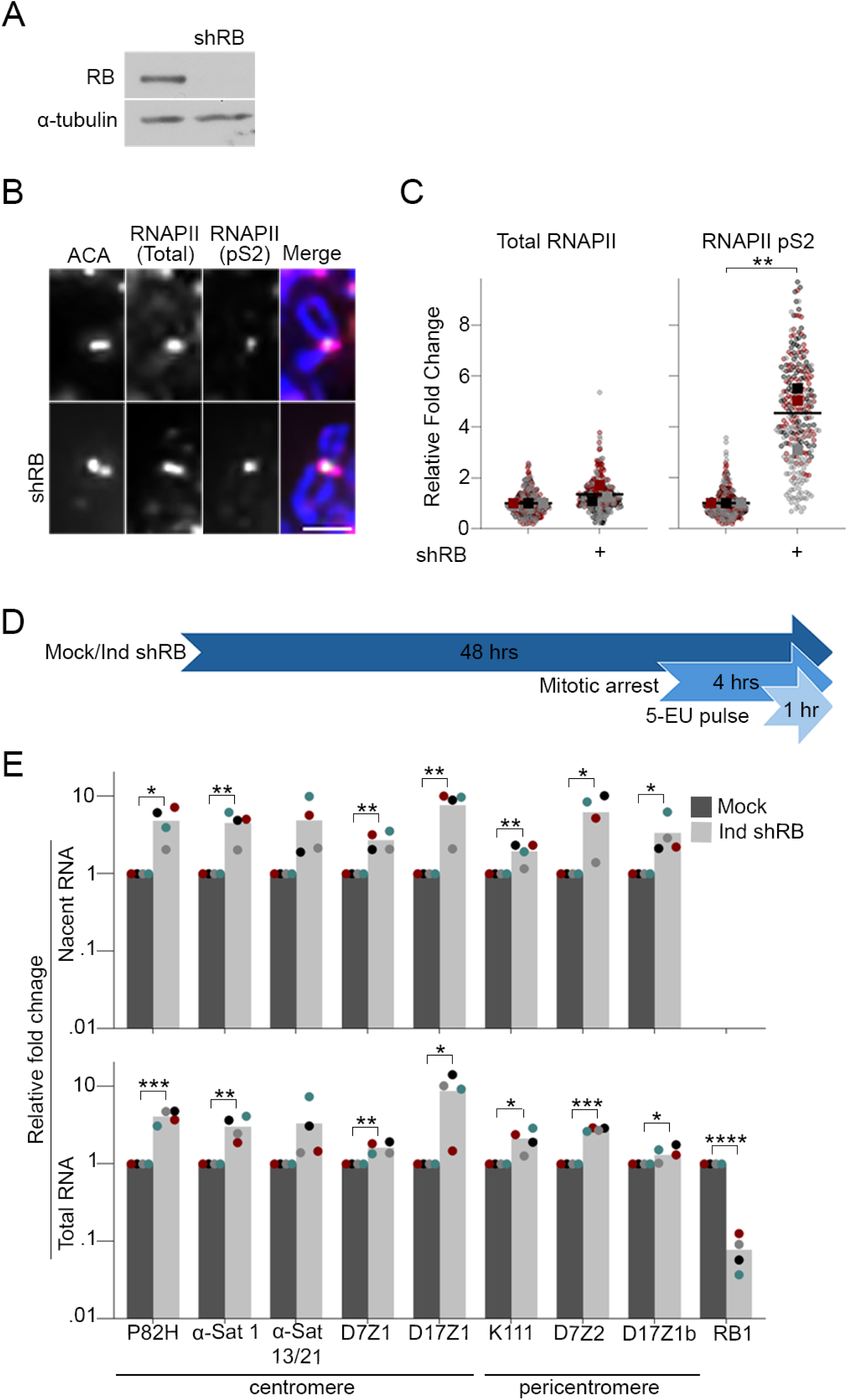
RB depletion increases RNAPII activity and transcription at the centromere. A) RB depletion via shRNA (shRB) in hTERT-RPE-1 cells was verified by Western blot. B) Representative images of mitotic spreads stained for total RNAPII (pink), active RNAPII (pS2, green), ACA (red), and DNA (DAPI, blue). Scale bar = 2μM. C) Quantification of RNAPII and RNAPIIpS2 signal intensity across ACA-labelled centromeres. A minimum of 90 kinetochore pairs were measured (3/cell for 30 cells), for each of 3 biological replicates. D) Schematic representation of protocol to label nascent mitotic RNA. E) qPCR analysis of nascent (EU-labelled RNAs) and total RNA in control and RB-depleted cells. Individual replicates are indicated by different colors, statistical analyses were performed between averages of biological replicates; *: p< 0.05, **: p<0.01; ***: p<0.001, ****: p<0.001.

To verify that RNA polymerase activity during mitosis corresponds with an increase in the synthesis of centromere transcripts, we first pulsed nocodazole-arrested mitotic cells with 5-ethynyl uridine (5-EU) for one hour. 5-EU is an analogue of uridine that is incorporated during RNA synthesis. Using click chemistry to link EU-labeled RNAs to biotin, followed by streptavidin bead pull down, newly synthesized mitotic RNAs were isolated from bulk cellular RNA (Figure 1D). Quantitative real-time PCR was used to perform comparative analysis of bulk and nascent centromere transcript levels from control and RB depleted mitotic hTERT-RPE-1 cells (Figure 1E). We find that mitotic cells lacking RB exhibit a significant increase in newly synthesized (nascent) centromere and pericentromere transcripts. A similar increase in transcript level is seen regardless of whether the sequence of the transcript analyzed is unique to a single chromosome (i.e. D7Z1/2 and D17Z1/b: centromere/pericentromere of chromosome 7 and 17, respectively) or common to multiple chromosomes (i.e. P82H: centromere and K111: pericentromere), suggesting an underlying, wide-spread dysregulation of transcriptional control at mitotic centromeres and pericentromeres in cells lacking RB.

### RB loss promotes DNA damage and ATR activation at mitotic centromeres

Unzipping of the DNA double stranded helix while the RNA polymerase reads and transcribes the nascent RNA leads to the formation of a transient 3-stranded DNA-RNA hybrid structure known as an R loop (Hamperl & Cimprich, 2016; Santos-Pereira & Aguilera, 2015; Thomas *et al*, 1976). Although a normal consequence of RNA transcription, these hybrid structures are prone to both single stranded (ssDNA) and double stranded DNA (dsDNA) breaks (Aguilera & García-Muse, 2012; Cohen *et al*, 2018). Perturbations in the balance between formation and resolution of R-loops contribute to DNA damage and genomic instability (Costantino & Koshland, 2018). Given our observation that loss of RB increases mitotic centromere transcription (Figure 1), and increased transcription is known to enhance R-loop- induced DNA damage (Crossley *et al*, 2019), we hypothesized that mitotic cells lacking RB may exhibit increased DNA damage at centromeres. To examine this possibility, hTERT-RPE-1 cells with and without induced expression of an RB-targeting hairpin (shRB) were arrested in mitosis for 4 hours, fixed and stained for the canonical DNA damage marker γH2AX (Figure 2A), and number of damage foci per cell quantified. This analysis revealed that cells lacking RB exhibit an increase in the fraction of mitotic cells exhibiting DNA damage (classified as 5 or more γH2AX foci in an individual cell; Figure 2B). Complementary approaches to examine γH2AX foci on mitotic chromosome spreads from control and RB-depleted cells indicate that many of these damage foci are localized proximal to centromeres (Figure 2C & 2D). Importantly, the increase in mitotic DNA damage following RB depletion is dependent on ongoing transcription during mitosis, as treatment of mitotic cells with the RNAPII inhibitor α-amanitin (administered concurrent with nocodazole-induced arrest) restored the proportion of cells exhibiting DNA damage to that seen in control cells (Figure 2A and 2B).

**Figure 2:**
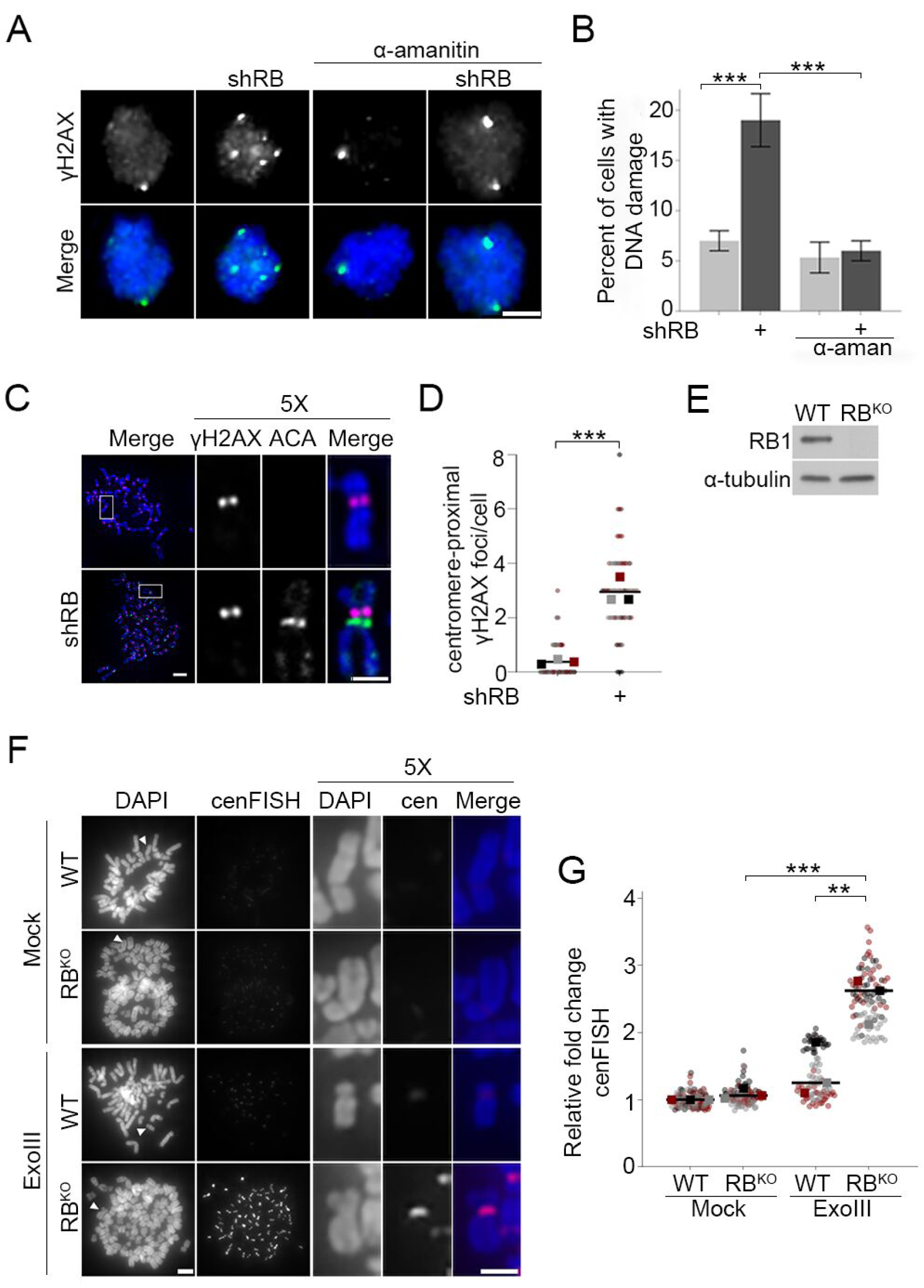
RB loss leads to centromere proximal DNA breaks. A & B) Representative images and quantification of γH2AX foci in mitotic hTERT-RPE-1 cells with or without induced shRNA-targeted depletion of RB (shRB) cells and/or treatment with α-amanitin (50μg/ml) during mitosis. Scale bar = 5μm, error bars represent standard deviation between biological replicates. C & D) Representative images and quantification of centromere-proximal γH2AX signal (green) in metaphase spreads co-stained for ACA (red) and DNA (DAPI, blue). A minimum of 90 kinetochore pairs were measured (3/cell for 30 cells), for each of 3 biological replicates. E) Western blot validation of RB loss in CRISPR-RB knockout hTERT-RPE cells (RB^KO^). F & G) Representative images and quantification of cenFISH probe labelling in WT and RB^KO^ cells following treatment (or not) with Exonuclease III. A minimum of 180 kinetochore pairs were measured (6/cell for 30 cells), for each of 3 biological replicates. In panels C& F white arrowheads or boxes indicate chromosomes represented in enlargements, scale bars are 2μm. Individual replicates are indicated by different colors, statistical analyses were performed between averages of biological replicates; **: p<0.01, ***: p<0.001.

While transcription-induced R-loops can lead to both single strand and double strand DNA breaks, γH2AX most efficiently labels double strand breaks (Rogakou *et al*, 1998; Sedelnikova *et al*, 2002). Therefore, to more comprehensively assess the extent to which RB-deficient cells acquire centromere and pericentromere nicks or breaks, we employed a fluorescent *in situ* hybridization (FISH) approach, termed exoFISH. In this assay, non-denaturing conditions limit centromere-targeting FISH probes to hybridize only when single-stranded DNA sequences are revealed following *in vitro* digestion with exonuclease III (ExoIII). Since ExoIII gains access via nicks in the DNA backbone, preexisting ss or dsDNA breaks enable ExoIII-dependent digestion (Rogers & Weiss, 1980; Saayman *et al*, 2023b) and a corresponding increase in FISH probe hybridization. Using a probe that specifically targets the centromere-localized CENP-B binding sites present on most chromosomes, we examined mitotic chromosome spreads from both control (WT) and RB knockout hTERT-RPE cells (RB^KO^) (Nicolay *et al*, 2015) for evidence of centromere-proximal breaks (Figure 2E). The specificity and sensitivity of this assay in revealing the presence of pre-existing DNA breaks are supported by negative controls in which both WT and RB^KO^ show low levels of centromere FISH probe hybridization in the absence of ExoIII treatment and positive controls in which WT and RB^KO^ cells pretreated with the DNA nicking enzyme Nt.BsmAI exhibit comparable levels of centromere FISH probe hybridization following ExoIII treatment (Figure 2F & 2G, Supplemental Figure 1A & 1B). Using this assay, we find that centromeres in RB^KO^ cells are more susceptible to exonuclease activity than control cells (Figure 2G). This increase in centromere FISH probe accessibility is indicative of increased damage at or near mitotic centromeres.

R-loops recruit and activate the DNA damage response element ATR (Kabeche *et al*., 2018). During mitosis, ATR activity at centromeres stimulates AurB kinase to promote kinetochore-microtubule turnover (Cimini *et al*, 2006). In agreement with published reports (Kabeche *et al*., 2018), we find that ATR is recruited to mitotic centromeres in both control and RB-depleted cells (Figure 3A & 3B). However, consistent with observations that mitotic cells lacking RB exhibit high levels of transcription-dependent DNA damage (Figure 2), ATR activity at the mitotic centromere (as measured by the presence of the autophosphorylation mark pATR-T1989) is increased following RB depletion (Figure 3A and Figure 3B). We additionally observe a comparable increase in recruitment of AurB kinase to mitotic centromeres (Supplemental Figure 2). Importantly, the increase in ATR activity and AurB localization are transcription-dependent, as treatment with α-amanitin reduces centromere-localized pATR and AurB to levels seen in control cells (Figure 3A and 3B, Supplemental Figure 2). AurB kinase promotes kinetochore-microtubule turnover, such that increased AurB localization at the mitotic centromere enhances microtubule release and results in mitotic errors (Cimini *et al*., 2006; Knowlton *et al*, 2006; Muñoz-Barrera & Monje-Casas, 2014) Consistent with this, we and others find that RB-deficient cells exhibit a high rate of mitotic segregation errors (Figure 3C & 3D; (Coschi *et al*, 2010; Hernando *et al*, 2004; Manning *et al*., 2010). In agreement with a model whereby increased transcription/high levels of R loops activate ATR and in turn enhance AurB activity, anaphase defects in RB-deficient cells are similarly suppressed by either RNAPII (α-amanitin) or ATR (VE-821) inhibition (Figure 3C & 3D).

**Figure 3:**
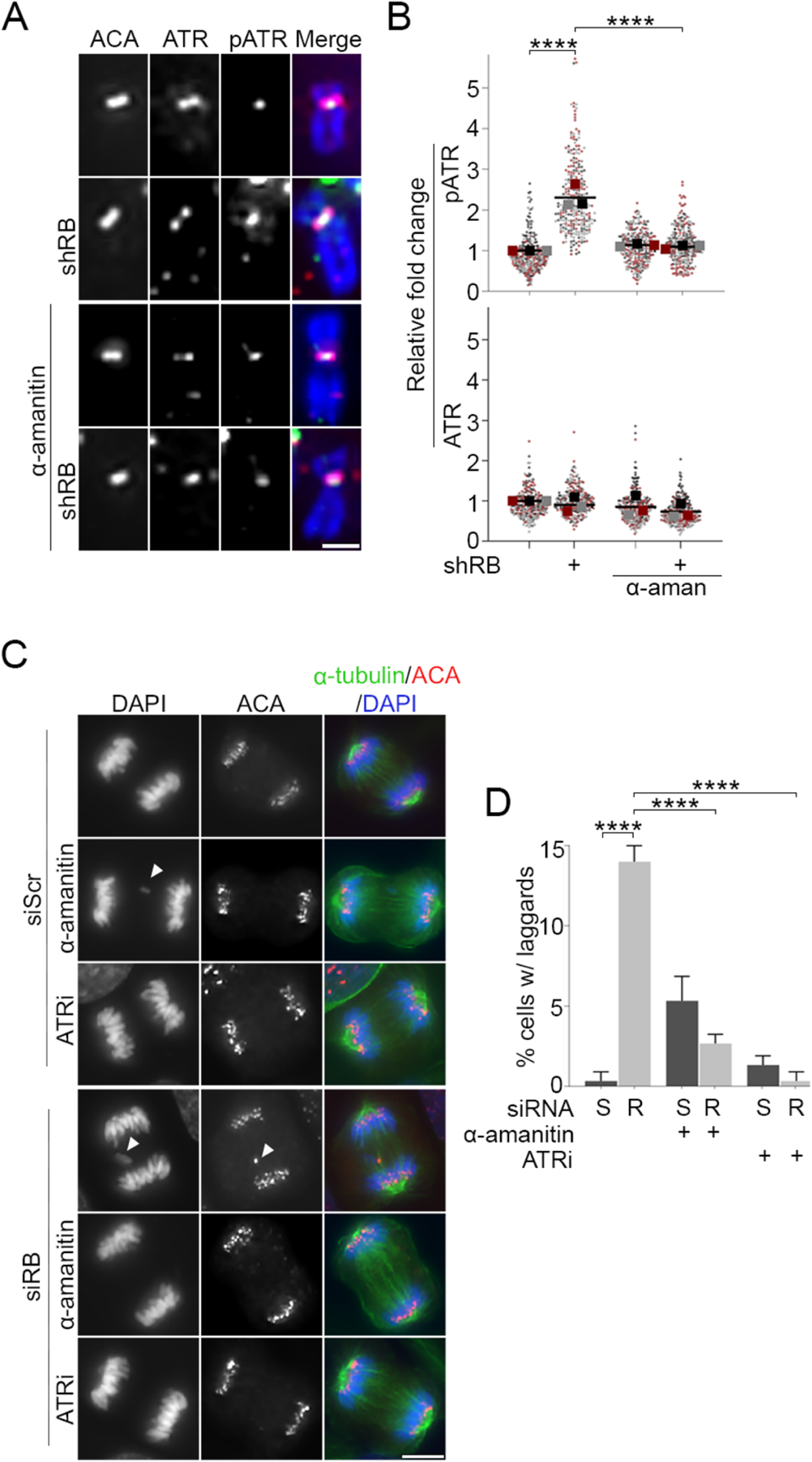
Transcription-dependent ATR activation compromises mitotic fidelity following loss of RB. A & B) Representative images and quantification of total (pink) and phosphorylated (T1989; green) ATR at mitotic ACA-labeled (red) centromeres in control and RB-depleted hTERT-RPE-1 cells. Cells were untreated or treated with the RNA polymerase II inhibitor α-amanitin (50μg/ml). A minimum of 90 kinetochore pairs were measured (3/cell for 30 cells), for each of 3 biological replicates. C & D) Representative images and quantification of anaphase defects in hTERT-RPE-1 cells treated with either a non-targeting control (S) or RB-specific (R) siRNA and subsequently treated with ATR inhibitor (VE-821, 10μM) for 1hr, α-amanitin (50μg/ml) for 4hrs. A minimum of 50 cells were scored per condition for each of 3 biological replicates. White arrowheads indicate lagging chromosomes, Scale bars are 2μm. Error bars represent standard deviation between biological replicates and statistical analyses were performed between averages of biological replicates, ****: p<0.0001.

### Epigenetic silencing of centromeres suppresses changes that result from RB loss

Pericentromeres are enriched with heterochromatin-promoting histone modifications including H3K9me3 and H4K20me3 (Jeppesen *et al*, 1992; Peters *et al*, 2001; Rea *et al*, 2000; Rice *et al*, 2003; Schotta *et al*., 2004). The RB protein has been shown to physically and functionally interact with a number of chromatin modifiers, including the H4K20 methyltransferase Suv420H2 (Gonzalo *et al*., 2005; Isaac *et al*, 2006; Siddiqui *et al*, 2007), such that loss of RB leads to decreased Suv420H2 and H4K20me3 enrichment at pericentromeres and telomeres (Gonzalo *et al*., 2005; Isaac *et al*., 2006). Given the transcriptionally repressive role of H4K20me3 (Gonzalo *et al*., 2005; Kourmouli *et al*, 2005; Schotta *et al*., 2004; Sullivan & Karpen, 2004) and our observation that RB loss leads to transcriptional upregulation of mitotic centromeres, we sought to explore whether loss of Suv420H2 enrichment may underlie centromere deregulation when RB is lost or depleted. To this end, we employed a previously established centromere-tethering system to anchor Suv420H2-GFP, via fusion to the DNA binding domain of centromere protein CENP-B, to centromeres (Herlihy *et al*., 2021). Using doxycycline-induced expression of a cen-Suv420H2-GFP fusion protein (Figure 4A), we first examined the impact of centromere-tethered Suv420-GFP (cenSuv) on mitotic centromere transcription in control and RB-depleted cells. As described above, nascent centromere RNAs were labelled with 5-EU, isolated, and quantified using qPCR. These data revealed that mitotic centromere transcription, which is increased following RB depletion alone, is reduced by concurrent tethering of Suv420H2 to the centromere (Figure 4B).

**Figure 4:**
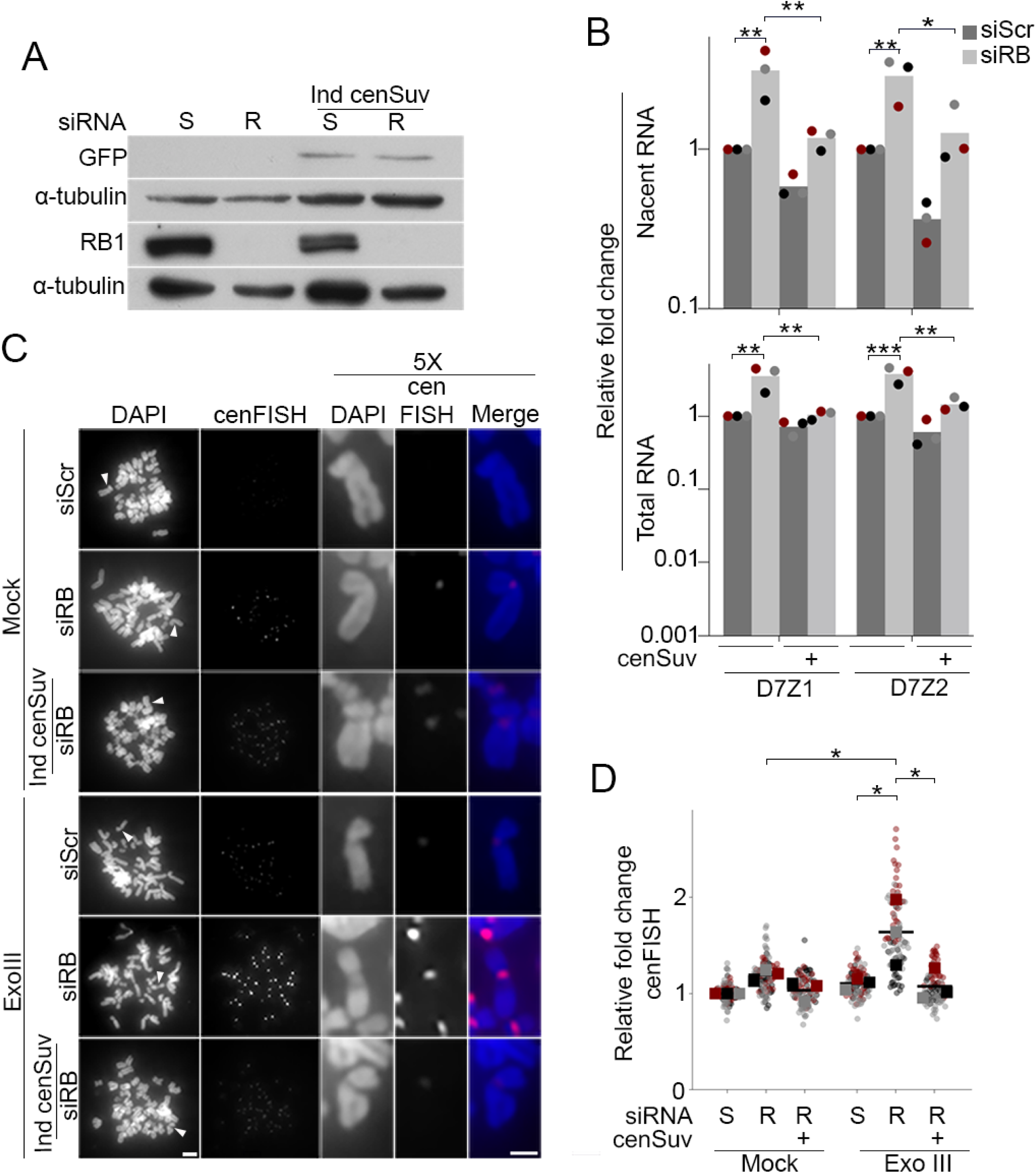
Decreasing mitotic centromere transcription reduces centromere breaks. A) Western blot analysis confirmation of RB knockdown and Suv420H2 overexpression in hTERT-RPE + inducible cen-Suv420H2-GFP cells treated with either a non-targeting control (S) or RB specific (R) siRNA, with or without induction of cen-Suv420H2-GFP expression. B) qPCR analysis of nascent (5-EU-labeled RNA) and total RNA transcribed from a representative centromere (D7Z1) and pericentromere (D7D2) in mitotic cells. C) Representative images and quantification of cenFISH signal in control and RB-depleted cells with or without Exonuclease III treatment. A minimum of 180 kinetochore pairs were measured (6/cell for 30 cells), for each of 3 biological replicates. Scale bars are 5μm for whole spread panels and 2μm for individual chromosome enlargement. Individual replicates are indicated by different colors, statistical analyses were performed between averages of biological replicates; *: p< 0.05, **: p<0.01; ***: p<0.001.

Treatment with α-amanitin indicates that increased centromere damage, ATR activation, and increased centromere AurB localization that occur following RB loss are dependent on RNA polymerase II activity (Figure 2, Figure 3). However, α-amanitin treatment alone cannot discern between a role for centromere transcription and more general transcriptional deregulation that may occur throughout the genome when RB activity is lost. Therefore, to more explicitly test the role of centromere regulation on these attributes in RB-deficient cells, we performed exoFISH, as described above, on control and siRB-depleted cells with and without expression of centromere-tethered Suv420H2-GFP. Upon induction of cen-Suv420H2-GFP the high level of ExoIII-dependent cenFISH signal seen in siRB depleted cells was reduced (Figure 4C & 4D; Supplemental Figure 3).

We next employed quantitative immunofluorescence to assess centromere levels of AurB kinase and anaphase defects following induced centromere tethering of Suv420h2-GFP in control and RB-depleted cells. Similar to RB depletion strategies described above (Supplemental Figure 2), siRB-treated cells exhibit an increase in the staining intensity of AurB at centromeres (Figure 5A & 5B) and an increase in anaphase defects compared to control cells treated with an si-Scramble sequence (Figure 5C & 5D). Consistent with suppression of both centromere transcription and centromere-proximal DNA damage described above, centromere tethering of Suv420H2-GFP in siRB cells reduced both centromere-localization of AurB (Figure 5A and 5B) and anaphase defects to levels comparable to that observed in cells treated with a scrambled siRNA control alone (Figure 5C & 5D).

**Figure 5:**
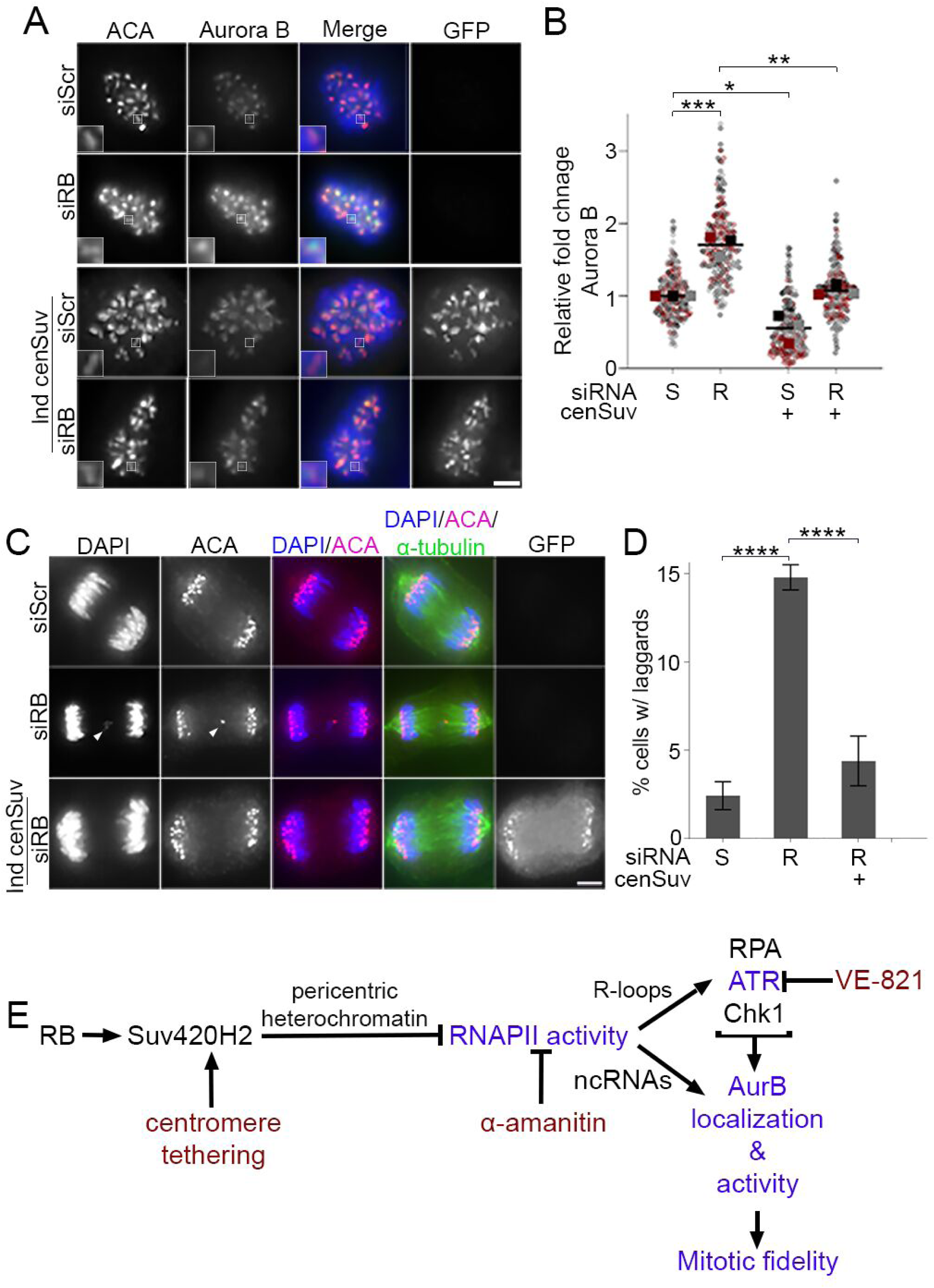
Suppression of transcription at the centromere limits Aurora B localization and reduces lagging chromosomes. A & B) Representative images and quantification of centromere-localized Aurora B (green) in hTERT-RPE + cen-Suv420H2-GFP cells treated with either a non-targeting control (S) or RB specific (R) siRNA, with or without induction of centromere tethered Suv420H2 and co-stained for ACA (red), GRF (white) and DNA (DAPI, blue). Insets are of individual kinetochore pairs at 3X magnification. A minimum of 90 kinetochore pairs were measured (3/cell for 30 cells), for each of 3 biological replicates. C) Representative images and quantification of anaphase defects in cells stained for ACA (red), tubulin (green), and DNA (DAPI, blue). A minimum of 50 cells were scored per condition for each of 3 biological replicates. Scale bars are 5μm. Error bars represent standard deviation between biological replicates. Individual replicates are indicated by different colors, statistical analyses were performed between averages of biological replicates; *: p< 0.05, **: p<0.01; ***: p<0.001, ****: p<0.0001. E) Model proposing how RB-dependent regulation of centromere transcription promotes mitotic fidelity. Dark blue text represents experimental readouts described in this study, red font represents experimental manipulations used to test/establish relationships.

## Discussion

Here we show that loss of the RB tumor suppressor permits increased RNA polymerase II activity at mitotic centromeres (Figure 1). The high temporal resolution afforded by first pulse labelling mitotic cells with 5-EU and then quantifying nascent RNA levels indicates that this feature of RB deficient cells corresponds with ongoing transcription at centromeres during mitosis (Figures 1 and 4). Our complementary approaches to either visualize the DNA damage marker γH2AX or alternatively to exploit ss and dsDNA breaks with exonuclease treatment reveal an increase in centromere-proximal DNA breaks following RB loss (Figures 2 and 4). We find that these breaks correspond with centromere-localized activation of ATR and AurB kinases (Figures 3, S2, 5). Consistent with previously described roles for ATR and AurB in the regulation of mitotic chromosome segregation, RB deficient cells exhibit an increase in anaphase defects (Figures 3 and 5). Importantly, inhibition of bulk mitotic transcription (RNA polymerase II inhibition with α-amanitin) or centromere-specific transcription (through centromere-tethering of the heterochromatin-promoting enzyme Suv420H2) limit mitotic DNA damage (Figures 2 and 4), reduce ATR and AurB kinase activity at centromeres (Figures 3 and 5), and restore mitotic fidelity in RB-deficient cells (Figures 3 and 5). These data support a model whereby high levels of centromere transcription renders cells lacking RB sensitive to DNA breaks and ATR activation, leading to increased AurB kinase localization and activity at centromeres. Together, these changes promote chromosome segregation errors and contribute to chromosome instability (Figure 5E).

### RB-dependent regulation of heterochromatic boundaries at centromeres is critical for mitotic fidelity

Proteomics analysis has indicated that the RB tumor suppressor protein physically interacts with over 300 proteins, a significant portion of which are epigenetic modifying enzymes (Sanidas *et al*., 2019). Functional studies of the RB interactome suggest that RB may serve as a scaffold-moderating where in the genome and when in the cell cycle distinct epigenetic modifiers interact with chromatin (reviewed in (Gonzalo & Blasco, 2005)). Thus, by impacting the recruitment of enzymes that place transcriptionally repressive marks at centromeres (i.e. H3K27me3 by EZH2 (Blais *et al*, 2007; Ishak *et al*, 2016), H3K9me3 by Suv39 (Nielsen *et al*, 2001; Vandel *et al*, 2001), and H4K20me3 by Suv420h2 (Gonzalo *et al*., 2005; Manning *et al*., 2014)) RB is poised to limit centromere and pericentromere transcription. Consistent with work showing that disruption of RB interaction with EZH2 in mouse models deregulate transcriptional repression (Ishak *et al*., 2016) we demonstrate here that loss of RB permits high levels of transcriptional activity at the centromeres in mitotic human cells and leads to defects in chromosome segregation during cell division. Furthermore, rescue experiments that exploit mitotic specific (via short term α-amanitin treatment) or centromere-localized (via centromere-tethered Suv420h2/centromere-specific H4K20me3 enrichment) transcriptional repression to limit segregation defects indicate that mitotic errors following RB loss result from deregulation of centromere transcripts late in the cell cycle, and may not otherwise be dependent on global changes in protein expression that occur from loss of RB-dependent repression of E2F transcription factors earlier in the cell cycle.

### Balancing centromere transcription for genome and chromosome stability

Prior studies have demonstrated that inhibition of RNAPII and the corresponding decrease in centromere transcription promote mitotic errors (Chen *et al*., 2021; McNulty *et al*., 2017; Rošić *et al*, 2014). These studies implicate centromere transcription in the establishment of open chromatin that is conducive for CENP-A deposition in preparation for the subsequent cell cycle (Bobkov *et al*., 2018), and the transcripts themselves in recruiting and tethering critical kinetochore components (Blower, 2016; Ferri *et al*., 2009; Jambhekar *et al*., 2014; Liu *et al*., 2015; McNulty *et al*., 2017; Quénet & Dalal, 2014). In the absence of centromere-derived transcripts the AurB kinase-containing chromosomal passenger complex (CPC) is not recruited to the kinetochore (Blower, 2016; Jambhekar *et al*., 2014). AurB kinase functions to destabilize kinetochore microtubules- a critical step in releasing improper attachments that form early in mitosis. In the absence of AurB activity, kinetochore-microtubule attachments are hyper stabilized and improper attachments persist, ultimately delaying biorientation and leading to segregation errors (Abe *et al*, 2016; Broad *et al*, 2020; Hauf *et al*, 2003; Huang *et al*, 2018).

In contrast to work showing that mitotic fidelity is sensitive to reduction of centromere transcription, de-repression and/or increased expression of repetitive sequences derived from or near centromeres, including LINE elements Satellite repeats, has been observed in various cancer contexts and is strongly correlated with poor patient prognosis (McNulty *et al*., 2017; Ting *et al*, 2011; Zhu *et al*, 2018; Zhu *et al*, 2011). While current studies have not yet discerned whether high levels of centromere transcription may be a driving force in tumorigenesis or merely a passenger that indicates widespread deregulation of transcriptional repression, recent molecular studies highlight several models whereby centromere transcription may directly impact genome stability and underlie cancer susceptibility (reviewed in (Petermann *et al*, 2022)). First, the process of transcription results in the generation of a DNA:RNA hybrid and displaced ssDNA structure known as an R loop. These structures are sensitive to both single strand and double strand DNA breaks and during S phase additionally pose an obstacle to replication that can promote replication stress and further DNA damage. To mitigate this risk, cells actively limit aberrant or excessive R-loop accumulation through the activity of DNA:RNA hybrid-specific endoribonuclease activity (Lockhart *et al*, 2019). Second, the presence of R loops activates the DNA damage response. During mitosis, this includes the recruitment of the ATR kinase to centromeres. ATR activity leads to local activation of AurB kinase (via an ATR-Chk1 axis) (Kabeche *et al*., 2018). As described above, AurB kinase functions to destabilize kinetochore microtubule attachments (Muñoz-Barrera & Monje-Casas, 2014) which can in turn compromise mitotic fidelity (Crowley *et al*, 2022; González-Loyola *et al*, 2015; Muñoz-Barrera & Monje-Casas, 2014). Activated oncogenes have been described to increase transcription and R loop formation, leading to replication stress and genomic instability (reviewed in (Petermann *et al*., 2022)). Our work additionally implicates the RB tumor suppressor, which is commonly lost or functionally inactivated across a broad range of cancer contexts (Burkhart & Sage, 2008), as a regulator of mitotic centromere transcription. We propose that, through recruitment of key epigenetic modifying enzymes, RB functions to establish and/or maintain a heterochromatic boundary at centromeres that in turn limits centromere transcription and AurB activity to promote mitotic fidelity. A number of groups have described that loss of RB compromises mitotic fidelity (Manning & Dyson, 2011). Data presented here are consistent with a model whereby, in the absence of RB, repressive marks near centromeres are reduced permitting excessive transcriptional activity. Aberrant R-loop formation and the corresponding recruitment of ATR kinase may then collaborate with transcript-dependent recruitment of AurB to destabilize kinetochore microtubules, concurrently leading to both centromere damage and whole chromosome segregation errors.

## Materials and Methods

### Cell culture

hTERT immortalized RPE-1 RB^KO^ (gift from the Dyson lab, Massachusetts General Hospital Cancer Center), RPE shRB (Zamalloa *et al*, 2023), and RPE cen-suv420h2 GFP (Herlihy *et al*., 2021) cells were grown in Dulbecco’s Modified Essential Medium (DMEM, Gibco) supplemented with 10% fetal bovine serum (Sigma) and 1% penicillin/streptomycin (Gibco). All cell lines were maintained at 37°C and 5% CO2. High resolution immunofluorescence imaging with DNA stain (DAPI, ThermoFisher) was used to monitor and confirm cell lines were free of *Mycoplasma* contamination. Over expression of cen-targeted suv420h2 GFP was achieved by treatment with 2µg/ml doxycycline for 16hrs. Inhibition of ataxia telangiectasia mutated and Rad3-related kinase (ATR) was achieved through treatment with VE-821 at a final concentration of 10µM (Sigma) for 1 hour. The inhibition of RNA polymerase II transcription was completed using a final concentration of 50ng/ml α-amanitin (Santa Cruz) for 4 hours. Depletion of RB1 was achieved through transient transfection with 50nM pool siRNAs (a pool of 4 siRNAs to RB1; siRNA-SMARTpool, Horzion Discovery) using RNAiMax transfection reagent (ThermoFisher) according to the manufacturer’s instructions. Transfection with a SMARTpool of four non-targeting siRNA sequences (Ambicon #AM4636) was used as a negative control for all depletions (Supplemental Table 1). Alternatively, cells were infected with a lentiviral construct containing a doxycycline inducible shRNA hairpin (tet-pLK0-Puro, Addgene #21915) for the targeted depletion of RB. Stable hairpin-expressing cells were selected with Puromycin for 10 days. Induced depletion was achieved by the addition of 2µg/ml doxycycline for a minimum of 48 hours.

### Immunoblotting

Cell extracts were prepared using 2x Laemmli buffer (BioRad) with β-Mercaptoethanol (Sigma Aldrich). Protein concentrations were normalized to total cell number and samples run on an SDS-PAGE gel. Proteins were transferred to PVDF membrane (Millipore) and blocked in 1xTBST supplemented with 5% milk. Antibodies were diluted 1:1000 in 1xTBST/5% milk: DM1Aα (α-tubulin, Santa Cruz), RB1 (4H1, Cell Signaling), GFP (D5.1, Cell Signaling) and incubated at 4°C. Membranes were washed in 1xTBST, incubated for in corresponding HRP-conjugated secondary antibody (GE Healthcare), and developed using ProSignal Pico (Prometheus).

### Immunofluorescence and Metaphase Spreads

Cultured cells were grown on coverslips, fixed, and stained for AurB and ACA as previously described in (Kleyman *et al*, 2014). To visualize anaphase defects and monitor DNA damage in nocodazole arrested prometaphase cells were fixed with 4% paraformaldehyde for 20 minutes, extracted with 0.2% TritionX-100 in PBS for 10 minutes, and blocked with TBS + 1% BSA. Primary antibodies, ACA (1:500, Antibodies inc), DM1A (1:1000 Santa Crux), or γH2AX Ser139 (1:1000, Cell Signaling) were diluted in TBS +1% BSA. Secondary antibodies were diluted in 1% BSA + 0.2mg/ml DAPI and coverslips were mounted onto slides using Prolong Antifade Gold (Molecular Probes). Mitotic cells were collected via shake-off after treatment with 100ng/ml nocodazole (Selleckchem) for 4 hours and metaphase spreads prepared as in (Keohane *et al*, 1996). Primary and secondary antibodies were diluted in KCM + 1% BSA. For staining of total (2B5,1:200 GeneTex) or phospho ATR (pT1989, 1:200 GeneTex) buffers were supplemented with a final concentration of 100nM Calyculin A (Millipore Sigma) (Kabeche *et al*., 2018). For total (F-12, Santa Cruz 1:200) or active RNAPII (pS2, Abcam 1:200) staining, buffers were supplemented with 100nM Calyculin A (Millipore Sigma), 40U/μL of RNasin (Promega) and kept on ice (Chan *et al*., 2012; Perea-Resa *et al*, 2020). Cells were then post fixed with 4% paraformaldehyde for 10 minutes prior to counter staining with TBS + 5% BSA + 0.2mg/ml DAPI for 30 minutes. Coverslips were mounted onto slides using Prolong Antifade Gold (Molecular Probes). Fixed cell images were captured using a Zyla sCMOS camera mounted on a Nikon Ti-E microscope, with a 60X Plan Apo oil immersion objective and 0.3 μm z-stacks. To assess centromeric protein levels, NIS-elements Advanced Research software was used to perform line scans in a single focal plane through individual ACA-stained kinetochore pairs where the area under the curve indicates the region of centromere/kinetochore-localized staining. γH2AX levels were assessed by counting the number of foci per cell. A cell was considered damaged if it had more than 5 foci per cell. For intensity measurements a minimum of 3 kinetochore pairs per 30 cells, per condition (90 kinetochore pairs/condition) were measured in each of 3 biological replicates. Anaphase defects were assessed in a minimum of 50 cells per condition for each of 3 biological replicates.

### exoFISH

exoFISH was performed as described in (Saayman *et al*, 2023a) with the following modifications. Cells were first prepared and spread onto coverslips as in (Ganem *et al*, 2009) Slides were then dried overnight at room temperature in the dark. The next day the slides were rehydrated in 1X PBS, treated with 0.5mg/ml RNaseA (NEB) for 10 minutes at 37°C in a humid chamber, washed with 1X PBS, then treated or not with 1 unit of Nt.BsmAI (NEB) for 2 hours at 37°C in a humid chamber. Slides were washed with 1X and treated or not with 200 mU/μL ExoIII diluted in 1X buffer supplied by the manufacturer for 1 hour then dehydrated overnight. To visualize breaks at the centromere slides were incubated in 0.5µM CENPB-binding site specific cenFISH probe (PNA Bio) for 3 hours at room temperature. Following incubation in hybridization wash buffers and subsequent dehydration, cover glass was then mounted onto slides using Prolong Antifade Gold. To quantify cenFISH signal, intensity values were measured within six 12x12 pixel boxes placed at centromeres and summed per cell for 30 cells per condition for each of 3 biological replicates. Fold change for each condition was calculated by normalizing to the RPE-1 without exoIII condition for that replicate.

Experimental data were analyzed with a Student t-test or one-way ANOVA where appropriate. Individual measurements from experiments where multiple measurements were made per replicate are represented as SuperPlots, with individual replicates color-coded. Per-replicate averages and standard deviation between biological replicates are superimposed. All error bars represent standard deviation between biological replicates and statistically significant differences are labeled with *: p<0.05, **: p<0.01, ***: p<0.001, and **** p<0.0001.

### Quantification of Nascent and total RNA levels

For quantification of nascent and total RNA levels mitotic cells were first isolated via mitotic shake off following incubation in 100 ng/ml nocodazole final concentration for 4 hours and subsequent treatment with 0.25µM 5-ethynyl uridine (5-EU, Vector Labs). Cells were processed according to the procedure outlined by the Click-iT™ Nascent RNA Capture Kit (Invitrogen). 2-5µg of RNA was removed prior to the remainder of the RNA undergoing the Click-iT™ reaction to be used for total RNA quantification and confirmation of RB knockdown. Total RNA was treated with DNaseI (NEB) before cDNA was synthesized from 1µg of total RNA using SuperScript IV Reverse Transcriptase (Invitrogen) according to the manufacturer’s instructions. cDNA synthesis for nascent RNA was performed according to the manufacturer’s instructions. Gene expression for centromeric and pericentromeric transcripts (Supplemental table 1) was determined using the ΔΔ cycle threshold method and normalized to GAPDH.

## Acknowledgements

The authors thank Nicole Hermance for technical support and critical evaluation of the manuscript. This work was supported by the American Cancer Society (RSG-21-066-01-CCG) to ALM.

## Author Contributions

ALM and EAC conceived and devised the study. EAC performed the experiments. ALM and EAC wrote the manuscript.

---

**Supplemental Figure 1:**
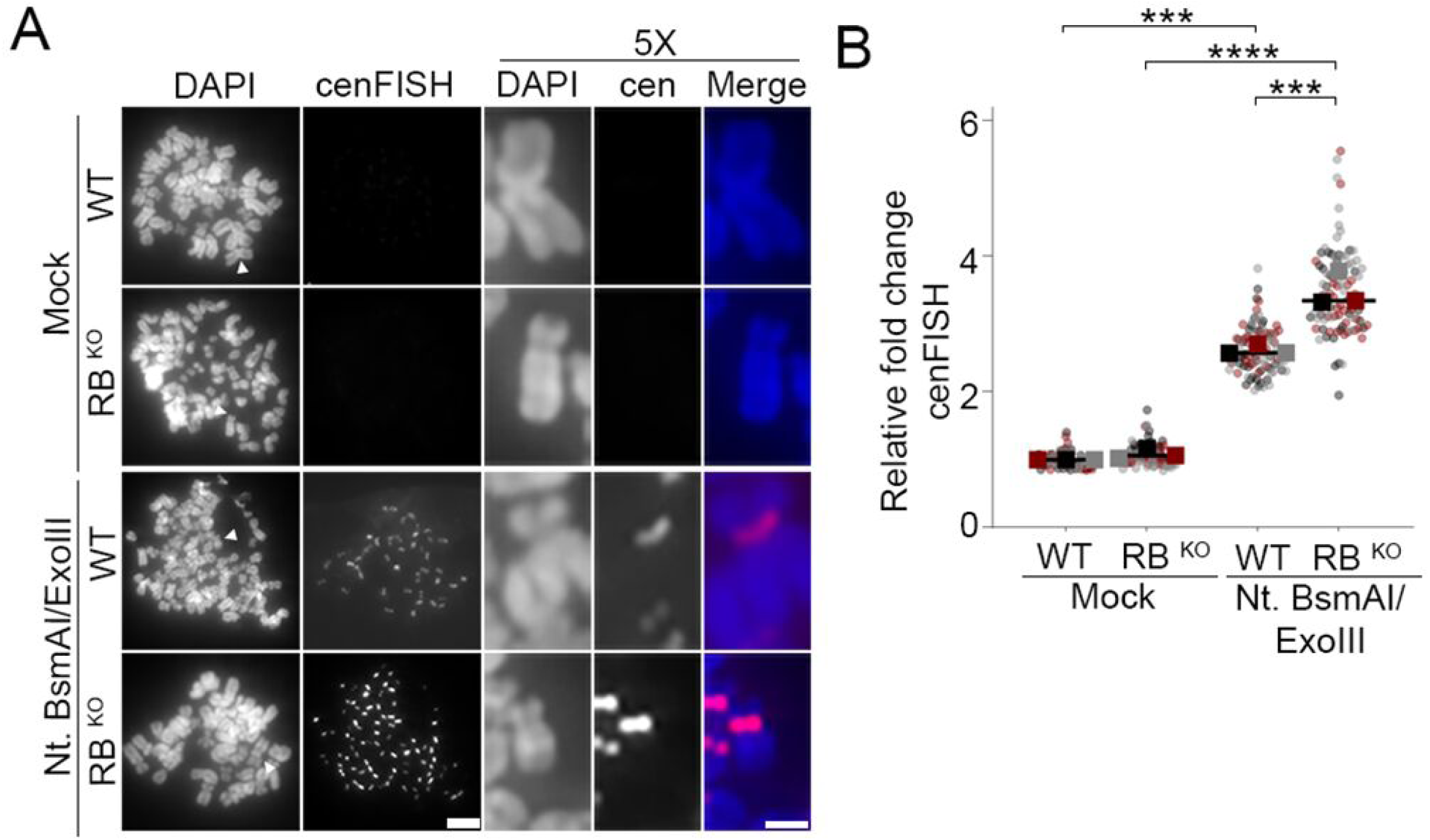
DNA breaks at the centromere can be detected through exoFISH. A & B) Representative images and quantification of centromeres in metaphase spreads of control (WT) and RB knockout hTERT-RPE-1 cells (RB^KO^) cells. Cells are labeled with a cenFISH probe following treatment, or not, with Nt.BsmAI and Exonuclease III. A minimum of 180 kinetochore pairs were measured (6/cell for 30 cells), for each of 3 biological replicates. Individual replicates are indicated by different colors, statistical analyses were performed between averages of biological replicates; *** p<0.001, ****: p<0.0001.

**Supplemental Figure 2:**
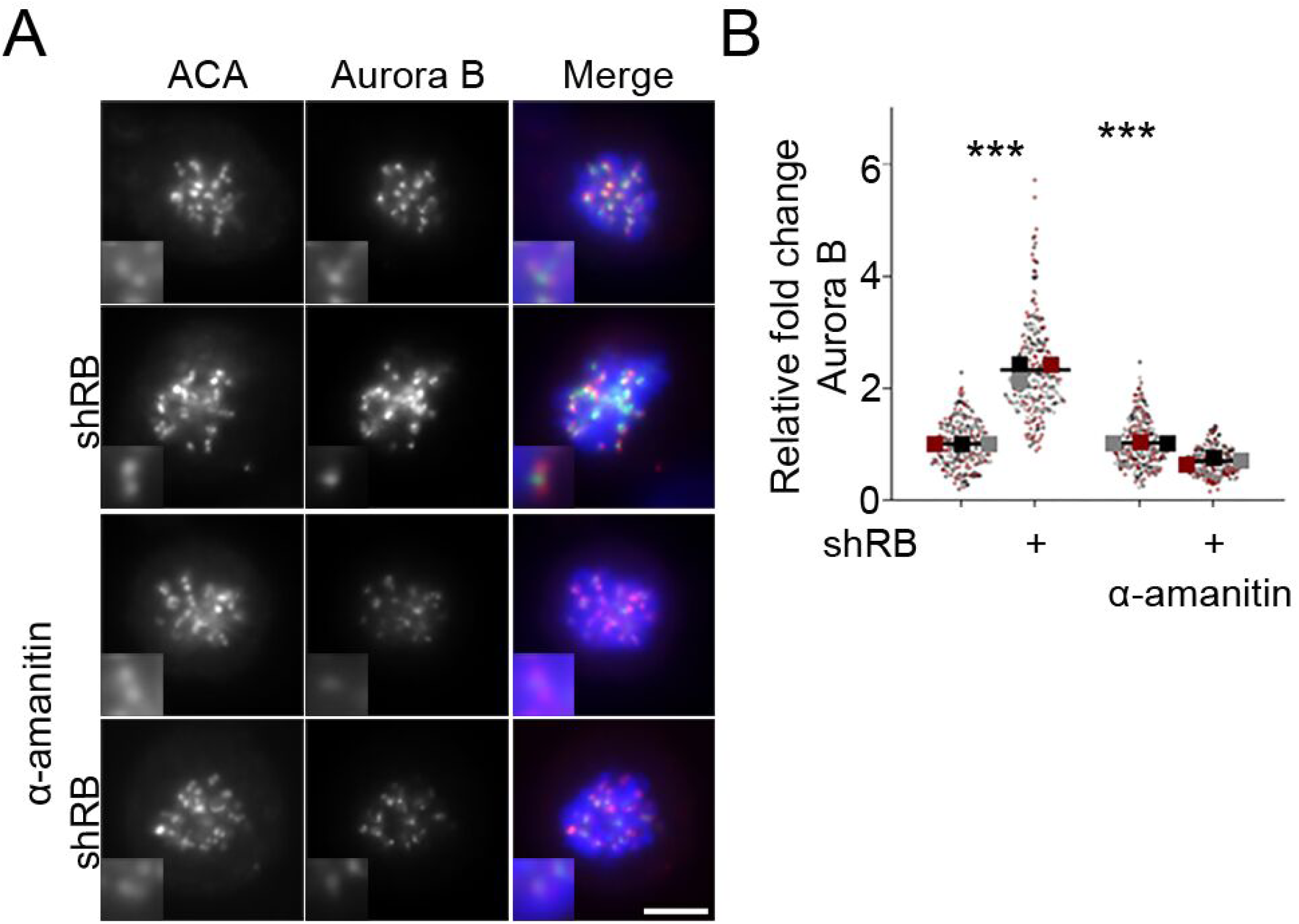
Aurora B localization is increased following RB loss. A & B) Representative images and quantification of centromere-localized Aurora B (green) in hTERT-RPE-1 cells co-stained for ACA (red), and DNA (DAPI, blue) following shRNA targeted depletion of RB (shRB) and/or treatment with α-amanitin (50μg/ml). Insets are of individual kinetochore pairs at 3X magnification. A minimum of 90 kinetochore pairs were measured (3/cell for 30 cells), for each of 3 biological replicates. Scale bar is 5μm. Individual replicates are indicated by different colors, statistical analyses were performed between averages of biological replicates; ***: p<0.001.

**Supplemental Figure 3:**
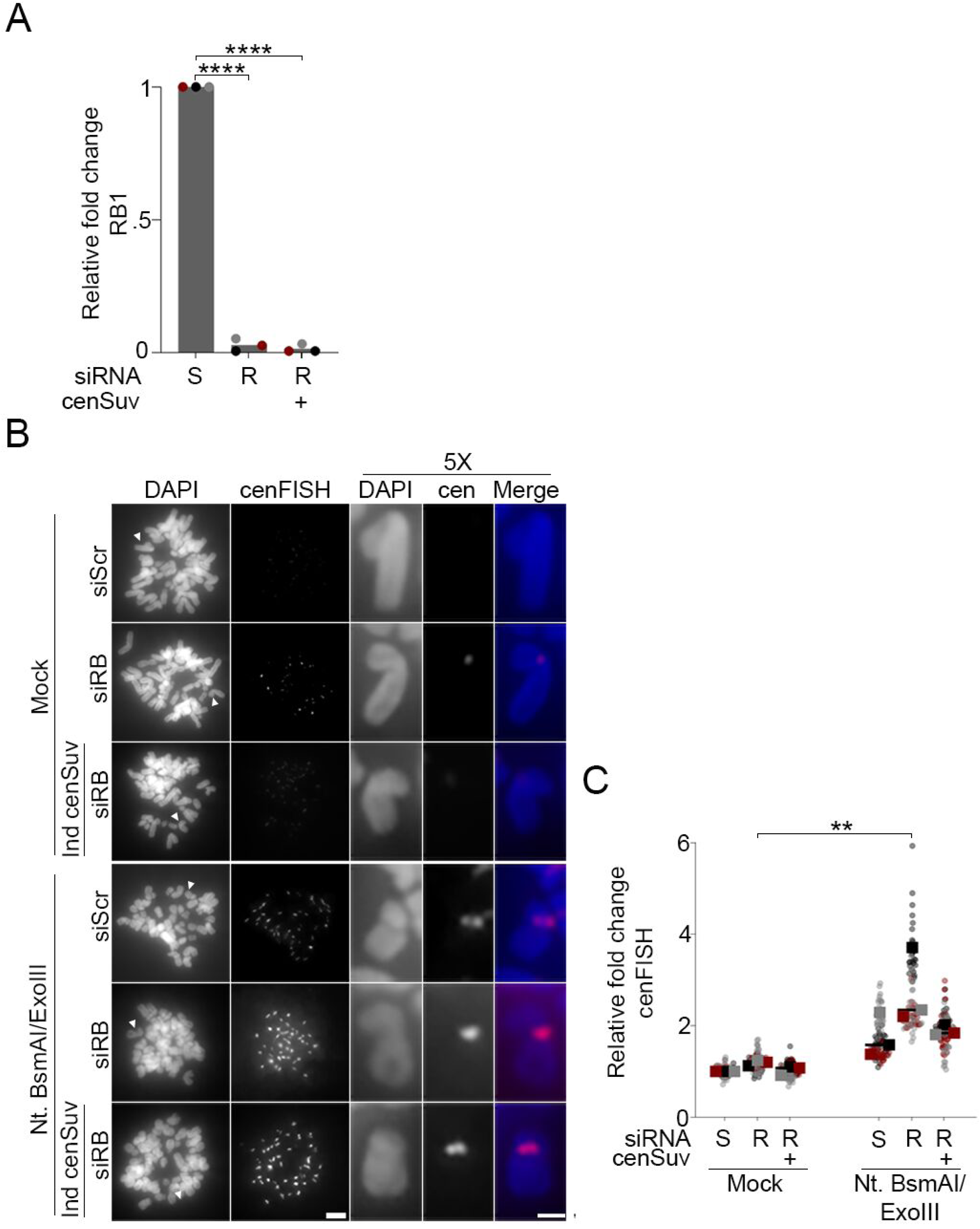
Validation of centromeric DNA break detection using exoFISH in cen-Suv420H2-GFP cells. A) qPCR analysis to validate RB depletion in hTERT-RPE + cen-Suv420-GFP cells treated with either a non-targeting control (S) or RB specific (R) siRNA, with or without induction of centromere tethered Suv420H2 expression. B & C) Representative images and quantification of centromeres in metaphase spreads. Cells are labeled with a cenFISH probe following treatment, or not, with Nt.BsmAI and Exonuclease III. A minimum of 180 kinetochore pairs were measured (6/cell for 30 cells), for each of 3 biological replicates. Scale bar is 5μm for whole spread images and 2μm for individual chromosomes. Individual replicates are indicated by different colors, statistical analyses were performed between averages of biological replicates; ** p<0.01; ****: p<0.0001.

